# Engineering a mouse-adapted SADS-CoV and establishing a neonatal mouse model to study its infection

**DOI:** 10.1101/2025.06.02.657353

**Authors:** Hanyu Zhang, Jiaru Zhou, Pengfei Li, Mengdi Zhang, Ran Jing, Hongmei Zhu, Yifei Lang, Qigai He, Mengjia Zhang, Wentao Li

## Abstract

Swine acute diarrhea syndrome coronavirus (SADS-CoV) is an emerging bat-origin alphacoronavirus causing severe disease in neonatal piglets, with significant economic losses to the swine industry as a consequence. The virus exhibits a broad species tropism, infecting cells derived from pigs, humans and mice, highlighting its potential for cross-species transmission. Due to drawbacks associated with the use of young piglets, there is a need for an appropriate small animal model to study SADS-CoV biology. Here we established a mouse infection model based on a murinized mutant of the virus, mSADS-CoV, in which the ectodomain of the viral spike protein was replaced by that of the murine coronavirus mouse hepatitis virus (MHV). This chimeric virus, generated through targeted RNA recombination, replicated efficiently in murine cell cultures and exhibited an age-dependent infection in neonatal mice that was lethal in 2-day-old BALB/c mice affecting various organs, notably the intestine. We validated our infection model by successfully verifying the efficacy of the RNA-dependent RNA polymerase inhibitor remdesivir (RDV). The model will serve as a valuable tool for studying SADS-CoV pathogenesis and for elucidating the roles of host factors in viral replication as well as for preclinical evaluation of vaccine candidates and antiviral compounds targeting the viral replication machinery.

**Author Summary:** SADS-CoV poses a dual threat to the swine industry and public health because of its broad species tropism and potential for cross-species transmission. The emergence of other bat-derived coronaviruses, including severe acute respiratory syndrome coronavirus (SARS-CoV), SARS-CoV-2, and Middle East respiratory syndrome coronavirus (MERS-CoV), underscores the need for robust models to study these pathogens. The successful rescue of mSADS-CoV and the development of a mouse infection model represent significant advancements in SADS-CoV research. This model not only enables the evaluation of antiviral therapeutics such as RDV but also provides a powerful platform for investigating viral replication mechanisms and host-pathogen interactions, offering critical insights for pandemic preparedness.

## Introduction

SADS-CoV is a recently identified porcine enteric coronavirus that predominantly affects neonatal piglets. Clinically, SADS-CoV presents symptoms strikingly similar to those caused by other known porcine enteric coronaviruses, including acute diarrhea and vomiting [1, 2]. Since its initial identification in Guangdong, China, in 2017, SADS-CoV has re-emerged in multiple regions, including Guangdong, Fujian, and Guangxi, giving rise to significant economic losses to the swine industry [3–5]. Emerging evidence suggests that SADS-CoV is likely of bat origin [2]. Notably, other bat-derived coronaviruses such as SARS-CoV-2, SARS-CoV and MERS-CoV have previously caused global outbreaks, posing severe threats to public health and the global economy [6–8]. This raises concerns that SADS-CoV may also hold a risk to human health through infection of an intermediate host. Studies have demonstrated that SADS-CoV can infect human cell lines, including Huh7.5, Caco-2, and ST-INT, and can efficiently replicate in cells derived from various other species, such as pigs, chickens, and mice, highlighting its significant potential for cross-species transmission [9, 10]. To date, no effective therapeutic agents or vaccines have been developed for the prevention or treatment of SADS-CoV, underscoring the critical importance of research into its pathogenic mechanisms.

While porcine neonates serve as natural reservoirs for SADS-CoV, the substantial experimental costs, inconsistent reproducibility, and technical complexity associated with porcine infection models significantly limit their utility in laboratory investigations. Mouse models generally offer distinct advantages for experimentation. Unfortunately, infection of SADS-CoV in mice exhibits inefficient replication stability. Previous studies have demonstrated that immunocompetent C57BL/6J mice (6-8 weeks old) sustain subclinical infections following viral challenge only transiently [10]. Subsequent investigations revealed markedly attenuated viral replication capacity in both immunodeficient and immunocompetent mouse hosts [9]. These findings emphasize the critical need to develop optimized mouse infection models that account for host immune cell heterogeneity as observed in African swine fever virus-associated lymphopenia [11] to facilitate rigorous *in vivo* characterization of SADS-CoV pathogenesis and therapeutic interventions.

SADS-CoV is an enveloped, positive-sense RNA virus distinguished by typical surface spike (S) glycoprotein protrusions. The 27.2 kb genome encodes four principal structural proteins: the S glycoprotein and the envelope (E), membrane (M), and nucleocapsid (N) proteins [2]. Proteolytic processing divides the S glycoprotein into S1 and S2 subunits, with the S1 subunit mediating receptor recognition and binding, while host proteases critically regulate S2 subunit activation to facilitate viral-host membrane fusion, a mechanism evolutionarily conserved among coronaviruses [12]. This functional specialization establishes the S1 subunit as the critical determinant of viral tissue tropism and host range [13, 14]. Owing to the essential biological role of the S glycoprotein, targeted RNA recombination strategies predominantly utilize S gene replacement to achieve viral recombination and selection [15].

Capitalizing on this molecular feature, we engineered in the present study a chimeric recombinant SADS-CoV carrying MHV spikes through targeted RNA recombination, designated as mSADS-CoV. Upon intraperitoneal administration in neonatal BALB/c mice (2-day-old), mSADS-CoV demonstrated robust replication competence with subsequent production of infectious virions. To test the performance of our animal model, we evaluated the efficacy of the broad-spectrum antiviral agent RDV against mSADS-CoV infection. The results demonstrated that RDV effectively suppressed viral replication *in vivo* and mitigated pathological tissue damage in infected mice. Notably, the model confirmed the physiological expression of replicase-associated genes during mSADS-CoV infection. The established mSADS-CoV infection system provides a valuable platform for various purposes including screening of replication-targeting antiviral agents, functional validation of host factors mediating coronaviral replication, and preclinical evaluation of therapeutic candidates and vaccine prototypes against SADS-CoV infection.

## Results

### Rescue of mSADS-CoV via targeted RNA recombination technology

With the aim of developing a murine animal model for SADS-CoV we engineered a murinized mutant of the virus, mSADS-CoV, by replacing the ectodomain of the spike protein by that of MHV using the prototype targeted RNA recombination technology [16](Fig 1A). The first step in the construction process involved the generation of a so-called transfer vector consisting of a genomic SADS-CoV cDNA from which ORF1a and almost all ORF1b is lacking and in which the sequence encoding the S protein ectodomain is replaced by the corresponding sequence from MHV. Thus, the small 5’-terminal genomic fragment and the ORF1b-Spike-3′UTR fragment were amplified from SADS-CoV cDNA template and ligated, meanwhile introducing a T7 promoter at the 5′-terminus. The amplified fragments were subsequently cloned into the pUC57 plasmid vector, generating pUC57-SADS-ORF1ab-Spike-3′UTR (Fig 1A). The proper spike sequence within this plasmid was then replaced by the corresponding MHV spike gene sequence amplified from MHV genomic cDNA, resulting in the transfer vector pUC57-SADS-ORF1ab-MHV Spike-3′UTR (Fig 1A). This plasmid was linearized and *in vitro* transcribed using T7 RNA polymerase, and the resultant mRNA was electroporated into SADS-CoV-infected Vero-CCL81 cells, which were subsequently overlaid onto mouse LR7 cells. Distinct cytopathic effects were observed in the LR7 cells (Fig 1B) indicating that recombinant virus mSADS-CoV had been generated. The virus was purified by two rounds of plaque purification after which its genomic structure was verified by sequencing. It grew in LR7 cells to a titer exceeding 10^6^ TCID_50_/ml, its replication kinetics showing a typical growth pattern (Fig 1C). The identity of mSADS-CoV was further confirmed through indirect immunofluorescence (IFA) assays, wherein all infected cells exhibited positive reactivity to both anti-MHV spike antibodies and anti-SADS-CoV N protein antibodies (Fig 1D), thereby validating the chimeric virus’s purity and structural identity.

**Fig 1.**
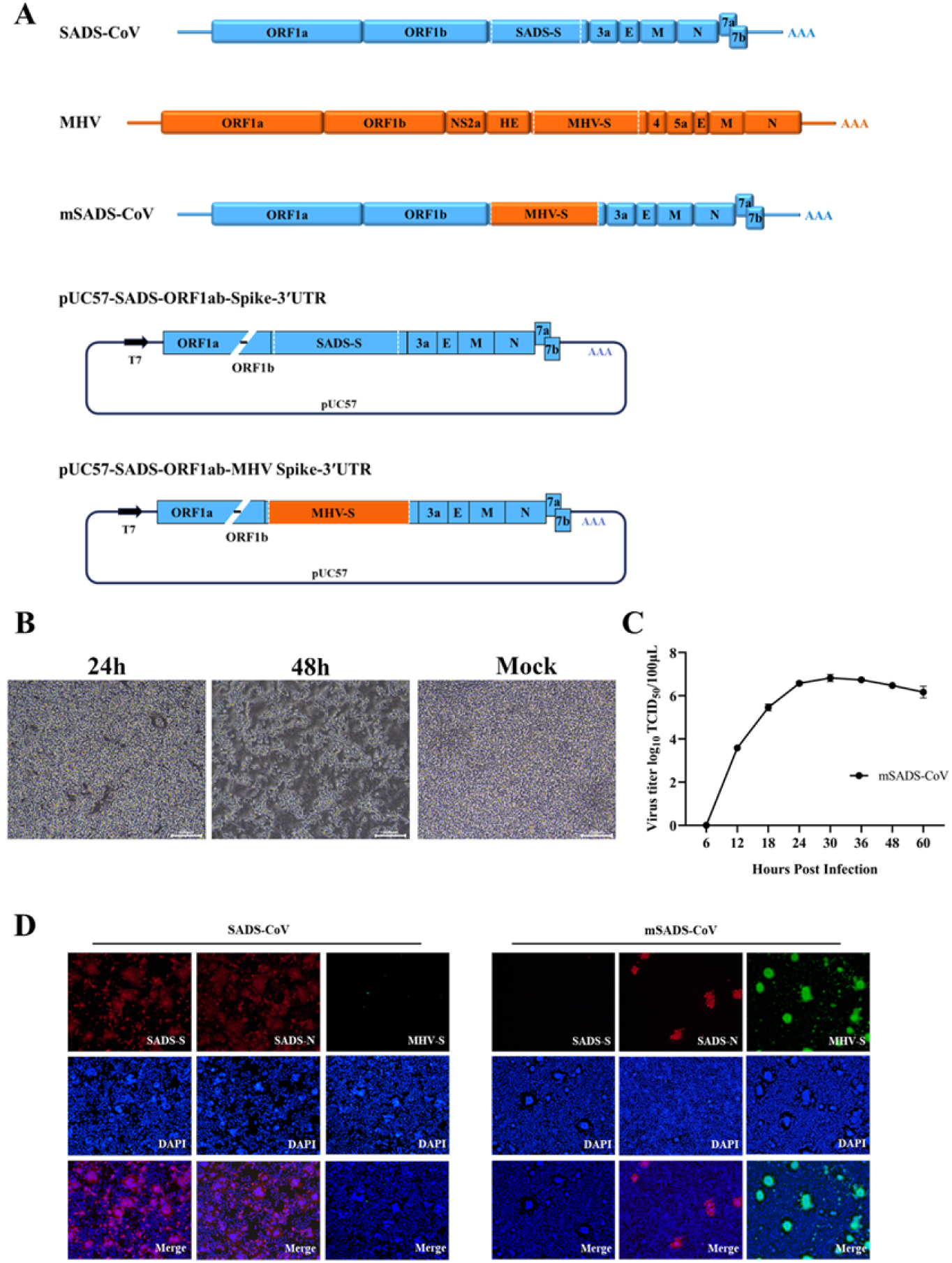
Rescue and validation of the recombinant virus mSADS-CoV. (A) Schematic representation of the generation of mSADS-CoV by homologous recombination of RNA, transcribed *in vitro* from pUC57-SADS-ORF1ab-MHV Spike-3′UTR and transfected into SADS-CoV infected Vero cells. (B) Cytopathic effects of recombinant mSADS-CoV in LR7 cells 24 and 48 hours post-infection. (C) Growth kinetics of mSADS-CoV in LR7 cells. Cells were infected with mSADS-CoV at an MOI of 0.01 and viral titers were determined by TCID_50_. (D) IFA of LR7 cells infected with mSADS-CoV using an anti-SADS-CoV S monoclonal antibody, an anti-MHV S protein antibody, and an anti-SADS-CoV N protein antibody. Vero CCL-81 cells infected with SADS-CoV were used as a control.

### Infection of two mouse strains with mSADS-CoV

To investigate the susceptibility of mice to mSADS-CoV, the virus was inoculated via intraperitoneal injection into young, 3 weeks old, mice from two different strains, BALB/c and C57BL/6. The clinical symptoms and body weight changes of the animals were monitored daily. Mice from neither group developed obvious clinical symptoms or mortalities. In BALB/c mice, no significant difference in average daily weight gain between the infected and control groups was observed; mSADS-CoV infection did not affect the animals’ weight gain (Figs 2A and 2C). In C57BL/6 mice, an initial more rapid increase in body weight in the infected animals relative to the controls was followed soon, from 3 to 4 days post inoculation, by a clear growth retardation (Fig 2B), which appeared to be significant as judged by a comparison of the average daily weight gain in each group (Fig 2C).

**Fig 2.**
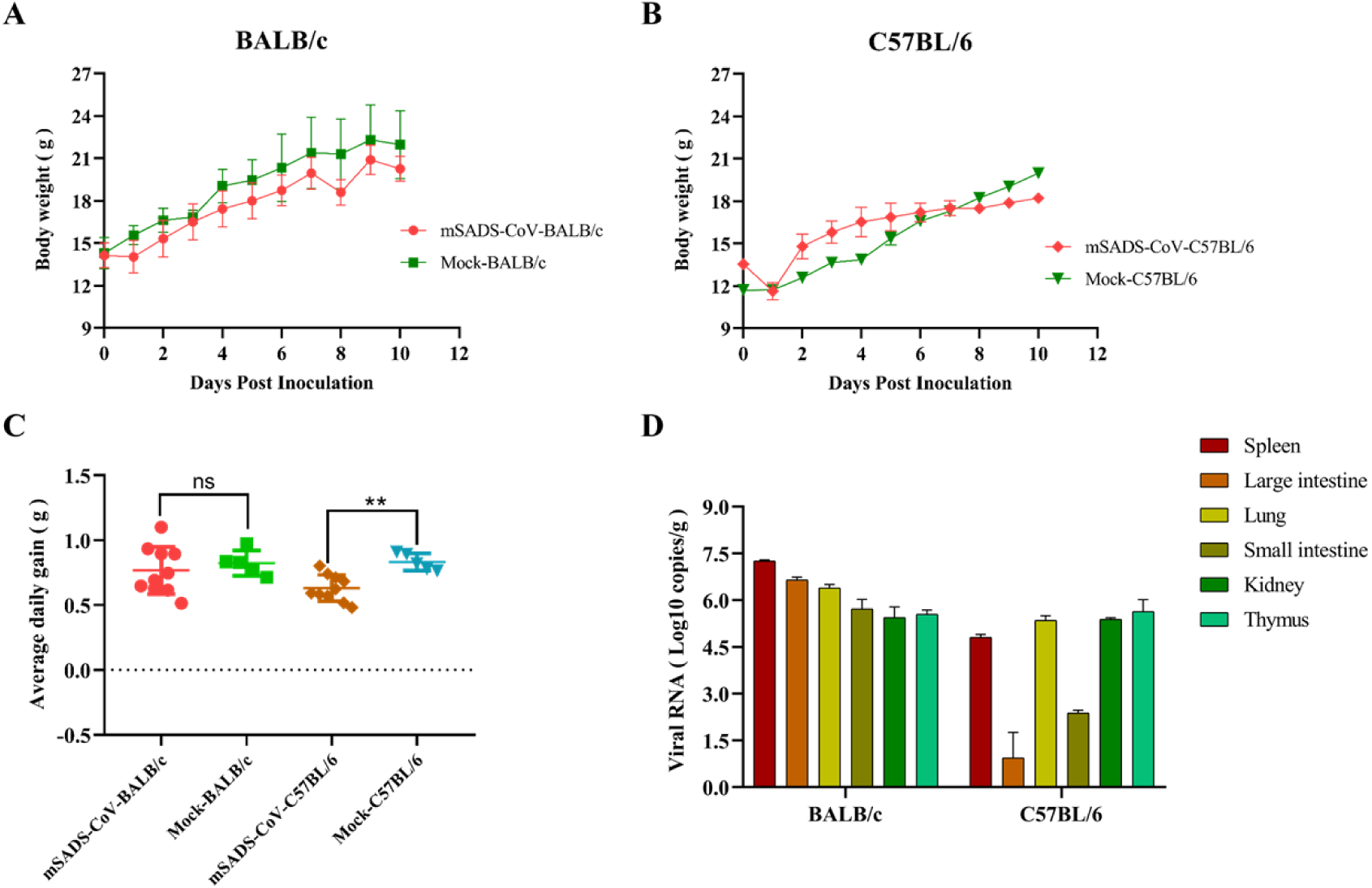
Infection of 3-week-old BALB/c and C57BL/6 mice with mSADS-CoV. (A) Body weight changes in BALB/c mice following the intraperitoneal administration of mSADS-CoV. (B) Body weight changes in C57BL/6 mice following the intraperitoneal administration of mSADS-CoV. (C) Average daily weight gain of BALB/c and C57BL/6 mice inoculated with mSADS-CoV via i.p. injection. (D) Viral loads in the spleen, large intestine, lung, small intestine, kidney, and thymus of BALB/c and C57BL/6 mice at 5 dpi by qRT-PCR.

Tissue samples from the mice were subjected to quantitative fluorescence PCR. A number of organs, including spleen, lungs, kidneys, thymus, large intestine and small intestine, tested positive in infected animals from both mouse strains. Viral loads in these tissues as measured at 5 days post-infection (dpi) are shown in Fig 2D. Except for the thymus, the viral loads in positive tissues of BALB/c mice were higher than those in C57BL/6 mice, with the viral loads particularly in intestines and lungs of C57BL/6 mice being significantly lower than those in other tissues. These results indicate that both BALB/c and C57BL/6 mice can be infected by mSADS-CoV through intraperitoneal injection, but that both strains exhibit subclinical infection. Based on the viral loads observed in the various organs, the virus replicates to higher titers in BALB/c compared to C57BL/6 mice.

### Infection of 2-day-old and 7-day-old mice with mSADS-CoV

In our search for a clinically more representative mouse model we conducted mSADS-CoV infection experiments using 2-day-old and 7-day-old BALB/c mice, taking SADS-CoV along in parallel for comparison. Inoculation of 2-day-old mice with mSADS-CoV resulted in the development of severe infection and disease. All animals died at days 5 or 6pi or had to be euthanized on day 6pi (Fig 3A). Some animals were already euthanized for necropsy at 3 dpi. In contrast, in the 7-day-old challenge group no obvious disease was observed and all mice survived. Body weight monitoring revealed that mSADS-CoV infection significantly inhibited normal weight gain in 2-day-old infected mice, whereas the average daily weight gain in the 7-day-old challenge group was not significantly different from that in the control group (Fig 3B). Quantitation of viral RNA by fluorescence PCR in several organs revealed that in the 2-day-old challenge group heart, liver, spleen, lungs, and intestines were all positive, with viral loads in each tissue exceeding 10^4.0^ copies/mg, reaching the highest titers of around 10^8.0^ copies/mg in the lungs and hearts at 5 dpi (Fig 3C). In the 7-day-old challenge group only the heart, lungs, spleen and intestines at 5 dpi, and the heart at 7 dpi were positive, the highest titers (in the lungs) being only 10^5.4^ copies/mg (Fig 3D). Necropsy of the mice from the 2-day-old challenge group at 5 dpi revealed that the walls of their large intestine were thinned, swollen and contained foamy contents (Fig 3E). To verify the presence of infectious viral particles in the positive tissues of the mice, tissue homogenates were prepared and their supernatants were inoculated onto LR7 cells. As shown by the IFA results (Fig 3G) all tested tissues appeared to contain infectious virus. These results indicate that mSADS-CoV can effectively replicate in various tissues of 2-day-old BALB/c mice, inhibiting weight gain and causing mortality, but that its infection efficacy decreases rapidly with age as no obvious clinical disease was observed when the animals were infected when 7 days old.

**Fig 3.**
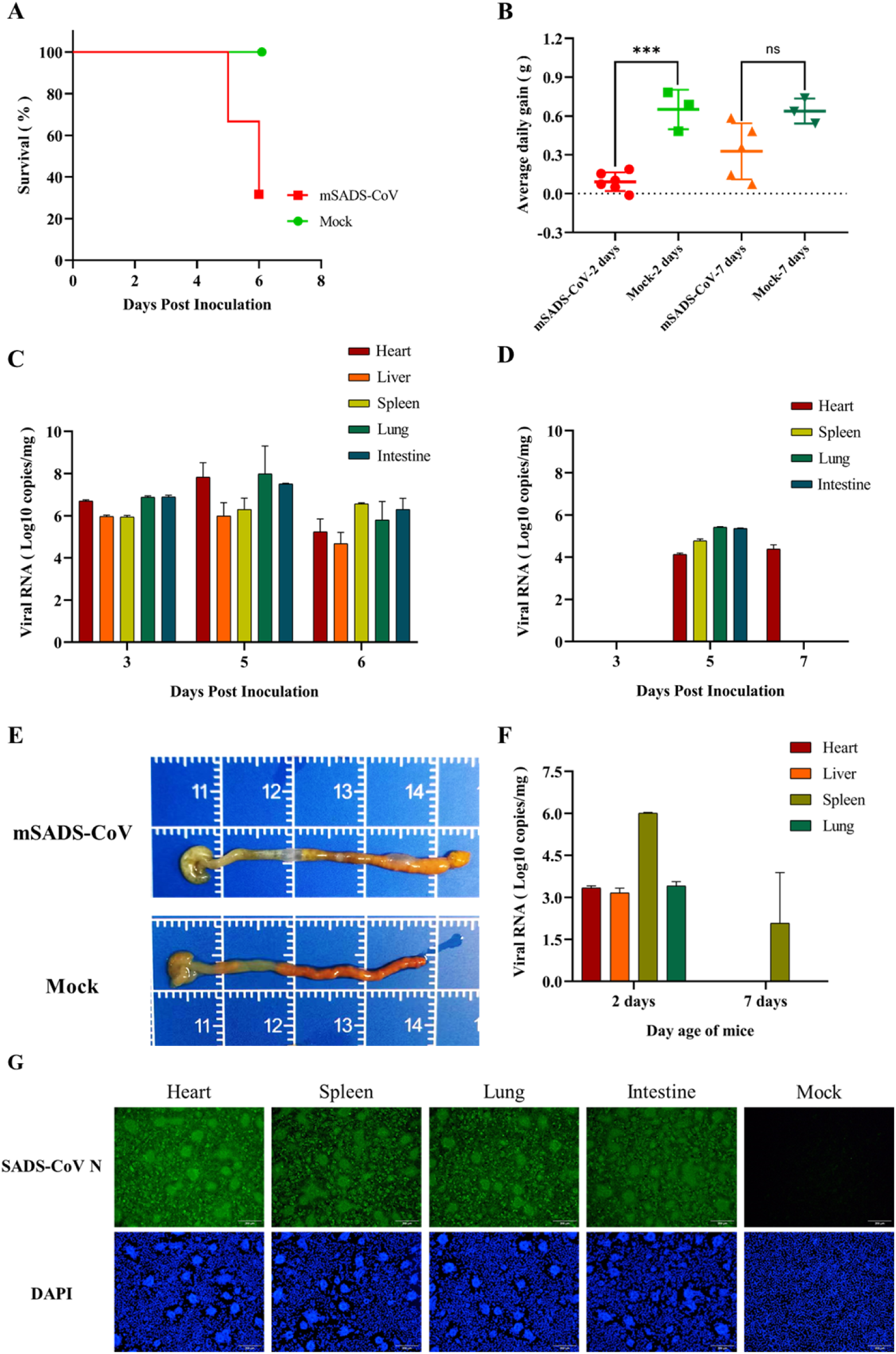
Infection of 2-day-old and 7-day-old BALB/c mice with mSADS-CoV. (A) Survival curve of 2-day-old BALB/c mice following i.p. injection of mSADS-CoV. (B) Average daily weight gain in 2-day-old versus 7-day-old BALB/c mice following i.p. injection of mSADS-CoV. (C) qRT-PCR determination of viral loads in the heart, liver, spleen, lung, and intestine of 2-day-old BALB/c mice after i.p. mSADS-CoV infection. (D) qRT-PCR determination of viral loads in the heart, liver, spleen, lung, and intestine of 7-day-old BALB/c mice after i.p. mSADS-CoV infection. (E) Pathological alterations in the large intestine of 2-day-old BALB/c mice at 5 dpi with mSADS-CoV. (F) Viral loads in heart, liver, spleen, and lung of 2-day-old and 7-day-old BALB/c mice at 3 dpi with SADS-CoV, as measured by qRT-PCR. The intestine tested negative in all cases. (G) IFA confirmation of successful isolation of tissue-derived virus using anti-SADS-CoV N protein antibody.

Intraperitoneal injection of BALB/c mice of the same ages and with the same dose of SADS-CoV did not elicit any significant clinical symptoms or weight loss. Yet, the animals had become infected as shown by the results of quantitative PCR analysis on some tissues collected at 3 dpi. In the 2-day-old challenge group, positive tissues included the heart, liver, kidney and, in particular, spleen, whereas in the 7-day-old challenge group only the spleen was positive, though with significantly lower titer (Fig 3F). These findings indicate that SADS-CoV can replicate - though poorly - in BALB/c mice, its infection exhibiting similar characteristics as mSADS-CoV, in particular with respect to their similar host age dependence. In both cases the virus is cleared quite rapidly by the host, making sustained infection difficult. We conclude that mSADS-CoV infection of 2-day-old BALB/c mice offers a suitable mouse infection model for SADS-CoV.

### Testing the mSADS-CoV mouse model by evaluating the antiviral efficacy of RDV

To test the applicability of the mSADS-CoV mouse infection model, we selected the broad-spectrum antiviral drug RDV for *in vivo* antiviral efficacy evaluation. We first checked whether RDV can inhibit mSADS-CoV infection *in vitro*. Thus, parallel cultures of LR7 cells were incubated for 2 hours with maintenance medium containing different concentrations of RDV. The culture supernatants were then replaced with culture media containing the same RDV concentrations and 0.1 MOI mSADS-CoV, and incubated for another 2 hours. The cells were then washed twice with PBS and incubated with maintenance medium containing again the same concentrations of RDV.

After 18 hours of mSADS-CoV infection, cell samples were fixed using 4% polyformaldehyde for IFA staining, while the supernatants were collected for subsequent TCID_50_ assay. The same protocol was used to verify the *in vitro* inhibitory effect of RDV on SADS-CoV, now in Vero CCL-81 cells. For both viruses the IFA analysis revealed that RDV can inhibit the replication of both SADS-CoV and mSADS-CoV in a dose-dependent manner (Figs 4A and 4B). The titers of SADS-CoV and mSADS-CoV in the supernatants decreased as RDV concentration increased, with a highly significant difference observed between RDV-treated groups and untreated controls. (Figs 4C and 4D).

**Fig 4.**
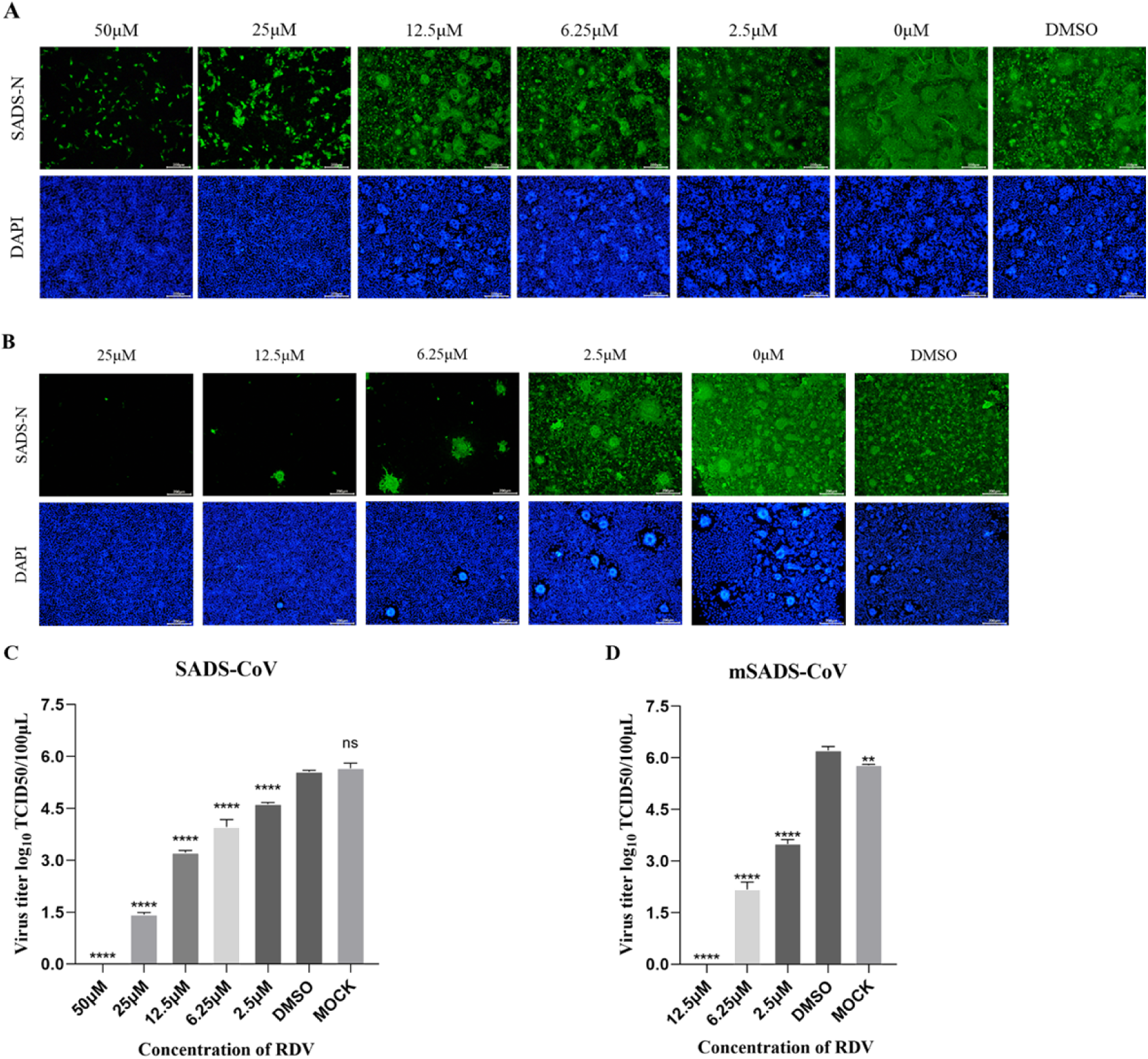
*In vitro* validation of RDV’s inhibitory efficacy against mSADS-CoV and SADS-CoV. (A) Determination of SADS-CoV infection in Vero CCL-81 cells treated with RDV at various concentrations by IFA staining. (B) Determination of mSADS-CoV infection in LR7 cells treated with RDV at various concentrations by IFA staining. (C) Viral titers of supernatants from SADS-CoV infected Vero CCL81 cells treated with different concentrations of RDV as expressed in log_10_ TCID_50_/100 μL. (D) Viral titers of supernatants from mSADS-CoV infected LR7 cells treated with different concentrations of RDV.

For the study in animals, 2-day-old SPF-grade BALB/c mice were randomly divided into three groups: an untreated and a RDV-treated infection group, and a control group. The animals in the infection groups were inoculated intraperitoneally with mSADS-CoV as before. The treatment group received subcutaneous injections of RDV at a dosage of 25 mg/kg/d every 24 hours starting 1 day before inoculation. Clinical observations and body weight monitoring revealed that all the mice in the RDV-treated infection group and the mock group survived throughout the observation period, whereas in the untreated infection group mice began to die from 4 dpi onwards, with a final mortality rate of 62.5% at 7 dpi, when the experiment was terminated (Fig 5A). While the RDV-treated animals grew at the same pace as those in the mock group, a severe growth retardation was observed in the untreated, infected mice as judged by the significantly lower average daily weight gain of this group (Fig 5B).

**Fig 5.**
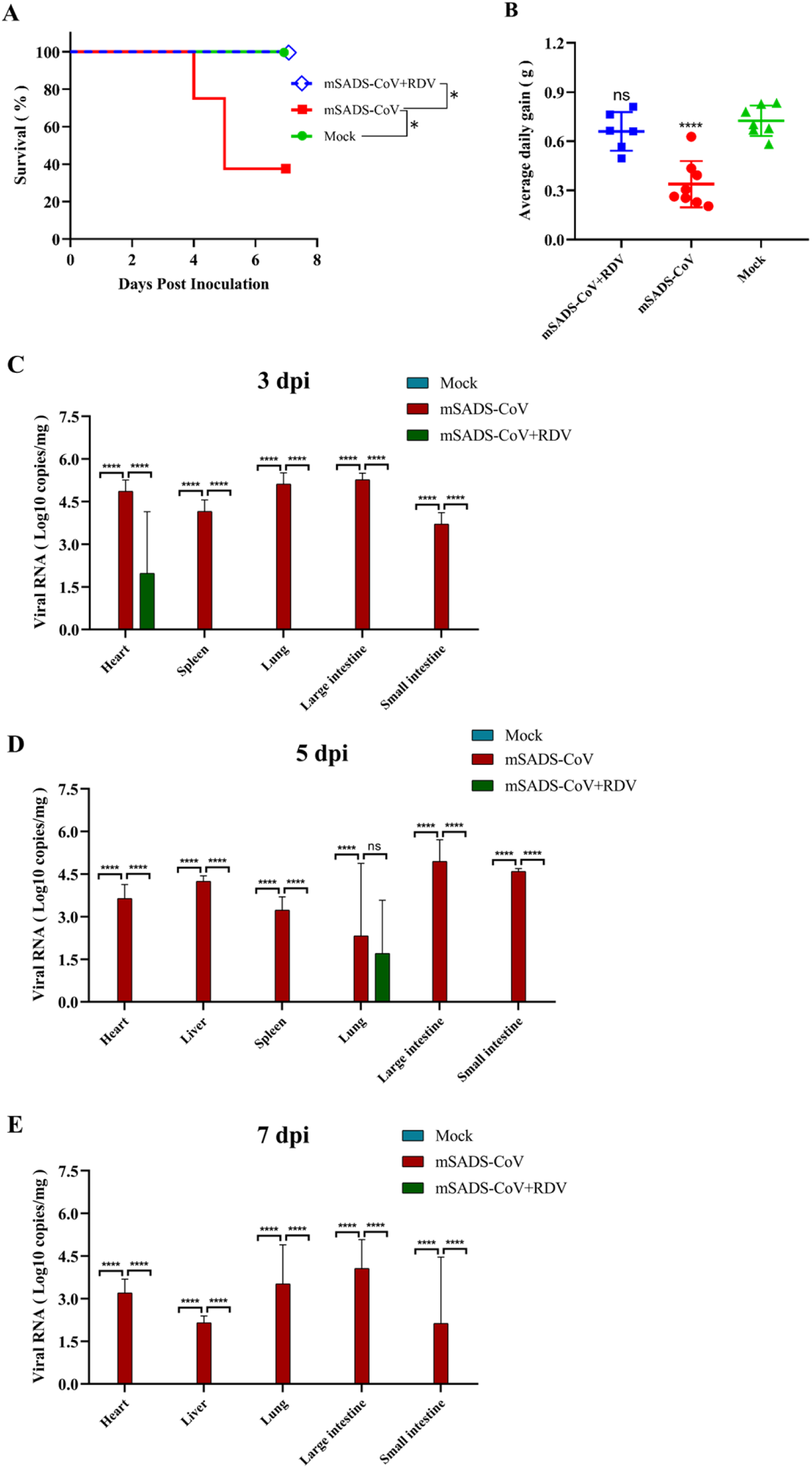
The effect of RDV on the infection by mSADS-CoV in 2-day-old BALB/c mice. (A) Survival curves of RDV-treated and untreated mSADS-CoV-infected mice and non-infected control animals. (B) Average daily weight gain of RDV-treated and untreated mSADS-CoV-infected mice and non-infected control animals. (C-E) Viral loads in heart, liver, spleen, lung, large intestine and small intestine of RDV-treated and untreated mSADS-CoV-infected mice and in non-infected control animals at (C) 3 dpi, (D) 5 dpi, and (E) 7 dpi.

Tissue samples collected from mice at 3, 5 and 7 dpi were subjected to quantitative fluorescence PCR to assess the impact of RDV on the viral loads in the tissues of the mSADS-CoV-infected mice. In the RDV-treated infection group viral genomic RNA was hardly detectible; only the heart at 3 dpi and the lungs at 5 dpi scored positive (Figs 5C-5E). In contrast, substantial viral loads were observed in all tested tissues from the untreated infection group, though the levels were somewhat lower than in the earlier experiment (Fig 3C).

The observations demonstrate that RDV significantly reduces viral replication in mSADS-CoV-infected mice, thereby reducing morbidity and mortality. Histopathological analysis of tissue samples from untreated mSADS-CoV infected animals taken at 5 dpi revealed distinct pathological alterations across multiple organs (Fig 6A): mild inflammatory cell infiltration was observed in cardiac tissue; hepatocytes exhibited variably sized lipid droplets; the splenic red pulp displayed marked expansion; pulmonary tissues exhibited alveolar epithelial hyperplasia, thickening of alveolar septa, and extensive inflammatory cell infiltration within the lung interstitium; and the large intestine presented a reduction in goblet cell numbers and disorganized epithelial cell architecture, accompanied by inflammatory cell infiltration in the mucosal layer. Of note, no significant pathological changes were detected in the small intestine. The RDV-treated infection group exhibited no notable histopathological alterations and was comparable to the control group. Immunohistochemical analysis further corroborated the efficacy of RDV in inhibiting mSADS-CoV infection in mice. Using a mouse monoclonal antibody targeting the SADS-CoV N protein, positive staining was observed in the alveolar walls of the lungs and the basal layers of the large and small intestines in the untreated mSADS-CoV group. In contrast, no clear staining was observed in the RDV-treated infected animals, as in the control group (Fig 6B).

**Fig 6.**
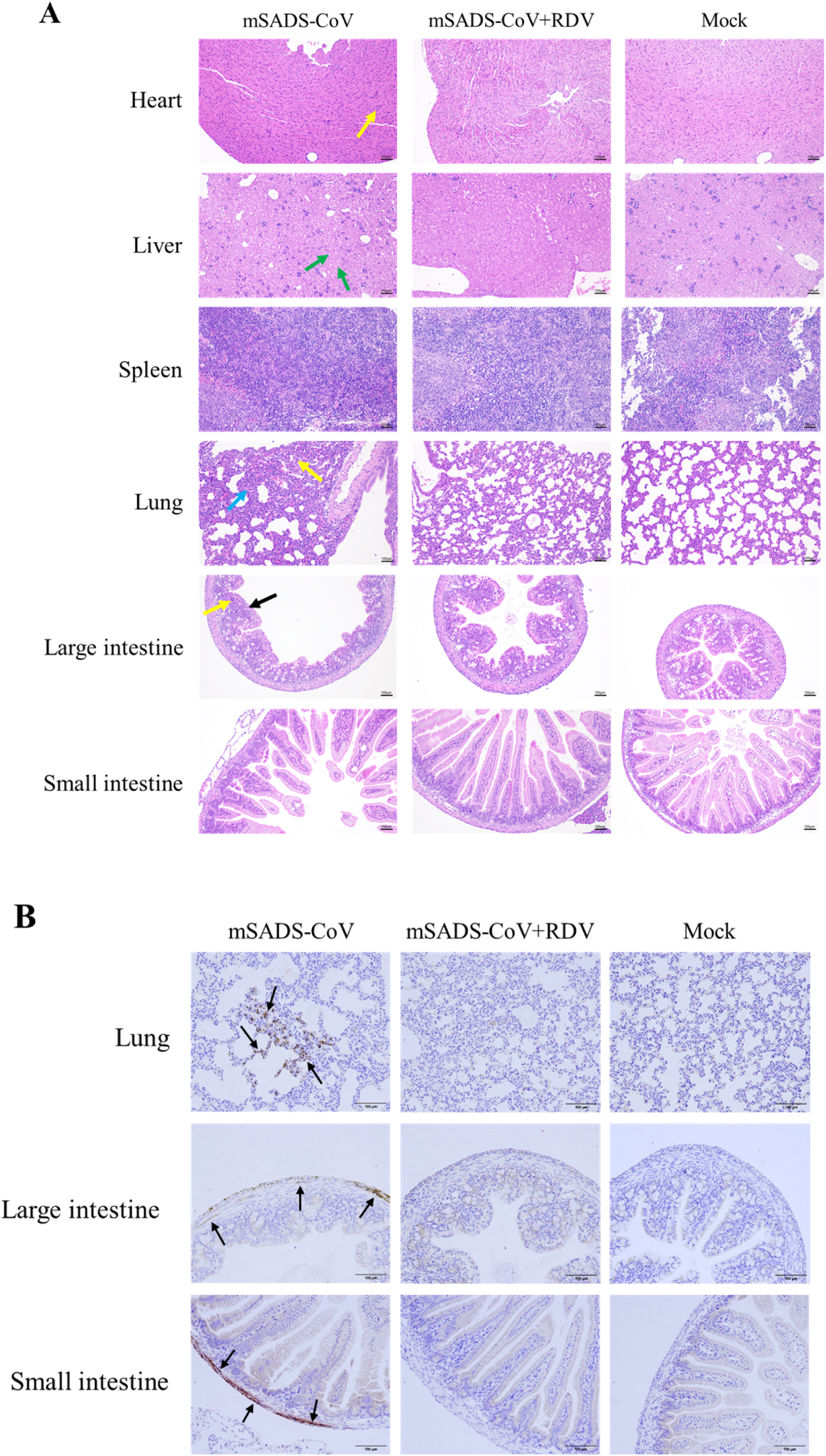
Pathological and immunohistochemical analysis of 2-day-old BALB/c mice infected with mSADS-CoV. (A) Hematoxylin and eosin (H&E) staining of different tissue sections from animals from the untreated infection group, the RSV-treated infection group and the control group at 5 dpi. *Yellow arrows*: inflammatory cell infiltration; *Green arrows*: polymorphic cytoplasmic lipid droplets; *Blue arrows*: concurrent alveolar epithelial hyperplasia and septal thickening; *Black arrows*: marked reduction of intestinal goblet cells. (B) Immunohistochemical (IHC) analysis of different tissue sections from animals from the untreated infection group, the RSV-treated infection group and the control group at 5 dpi. *Black arrows*: mSADS-CoV antigen distributed in the alveolar walls of the lungs and the basal layers of the large and small intestines (representative positive signals are indicated).

## Discussion

The emergence of several porcine enteric coronaviruses has reached a stage where these viruses are collectively responsible for considerable economic losses in global swine industries. Scientific evidence has convincingly demonstrated the distinctive identity of SADS-CoV compared to other prevalent porcine enteric pathogens such as porcine epidemic diarrhea virus (PEDV), porcine deltacoronavirus (PDCoV), and transmissible gastroenteritis virus (TGEV) [17, 18]. Immunologically this distinction directly correlates with the lack of cross-protective efficacy of existing commercial vaccines developed for these related viral pathogens against SADS-CoV infection. Even more distinctive is SADS-CoV’s demonstrated broad cellular tropism, which constitutes a substantial risk of interspecies transmission. Epidemiological evidence suggests SADS-CoV to have emerged from bats into rodents by cross-species transmission before spilling over into porcine hosts, its establishment in domestic swine populations creating substantial zoonotic risks through amplified human exposure opportunities[19, 20]. This transmission paradigm raises significant concerns regarding potential anthroponotic spread. Elucidating the pathogenic mechanisms underlying SADS-CoV infection therefore represents a critical research priority for public health security. Concurrent advancements in developing prophylactic and therapeutic countermeasures are urgently required to mitigate pandemic risks. The establishment of robust, reproducible animal infection models constitutes a fundamental prerequisite for investigating viral pathogenesis and for enabling the pre-evaluation of drug and vaccine candidates.

While neonatal piglets demonstrate susceptibility to SADS-CoV infection, the utilization of porcine models presents several methodological limitations, including experimental variability, technical complexity, prolonged reproductive cycles, and limited availability of genetic manipulation tools, thereby restricting their utility in molecular virology research. In contrast, mouse models offer distinct advantages, characterized by rapid reproductive turnover, comprehensive genetic toolkits, experimental tractability, and well-established knockout mouse repositories. The advanced development of mouse transgenic and gene-editing technologies has proven particularly valuable for investigating viral receptor usage, viral life cycle dynamics, and pathogenic mechanisms. Current research on SADS-CoV mouse models remains in its nascent stage, with inconsistent infection outcomes reported across studies. Experimental evidence indicates that SADS-CoV induces subclinical infection in 6-8 week-old C57BL/6J mice, with detectable viral replication in splenic tissue [10]. Other investigators, however, detected minimal viral replication in 10-week-old BALB/c immunocompetent and immunodeficient mouse models [21]. Chen et al. identified age-dependent susceptibility in both BALB/c and C57BL/6J mice, with neonatal mice (< 7 days) exhibiting highest vulnerability and showing progressive loss of susceptibility with increasing age, animals 3 to 4 weeks old appearing fully resistant [22]. Duan et al. successfully established infection in 7- and 14-day-old BALB/c mice via intracerebral inoculation [23]. However, our laboratory’s preliminary attempts to infect neonatal mice yielded inconsistent results, suggesting instability in this infection model. Furthermore, the limited blood-brain barrier permeability of most clinical antiviral compounds poses significant challenges for utilizing intracerebral inoculation models in antiviral drug screening protocols.

In order to improve or create a mouse infection model, one of the currently emerging approaches is by generating transgenic animals made (more) susceptible by the introduction of the cell entry receptor for the particular virus. Well-known examples in the field of coronaviruses are the mouse strains that express the human receptor for SARS and MERS coronaviruses, ACE2 [24, 25] and DPP4 [26], respectively. This approach obviously requires the relevant receptor to be known, which is not the case for SADS-CoV. As an alternative, rather than adapting the mouse to the virus, an inverse approach is feasible as well. In its classical format this may be achieved by passaging the virus serially through mice, thereby enabling the virus to evolutionary adapt to the murine host. A more directed approach to achieve such adaptation involves the retargeting of the virus by genetic engineering. The ability to do so was originally demonstrated some 25 years ago; by replacing the spike protein’s ectodomain from one coronavirus by that of another, the resulting chimeric virus adopted the tropism of the latter. Thus, MHV was retargeted to feline cells using a feline coronavirus S sequence [16], the feline coronavirus was inversely retargeted to mouse cells [27] and an avian coronavirus similarly to mouse cells [28].

A convenient way to generate such chimeric coronaviruses is by targeted RNA recombination technology [15]. As also demonstrated in the present study, this method enables the selection of the recombinant virus simply by growth on cells of the intended target species, in our case mouse cells. As MHV naturally infects mouse hosts, exhibiting high infectivity and pathogenicity characterized by hepatitis, enteritis, thymic atrophy, and induction of autoimmune responses [29, 30], the chimeric mSADS-CoV generated in this study theoretically possesses enhanced mouse infectivity compared to wild-type SADS-CoV, a hypothesis substantiated by our findings. The successful application of S protein ectodomain replacement in rescuing multiple coronaviruses [16, 28, 31–33] establishes its utility as a versatile platform for coronavirus research. This methodology can be effectively employed to generate chimeric recombinants of diverse coronaviruses, particularly facilitating the development of mouse-adapted viral variants. Such advancements provide a novel methodological framework for establishing mouse infection models across various coronavirus species, thereby enhancing our capacity to study coronavirus pathogenesis and host adaptation mechanisms.

Our observations demonstrated that SADS-CoV caused subclinical infection in neonatal BALB/c mice, exhibiting limited replicative capacity in 2-day-old, less in 7-day-old animals, followed by rapid viral clearance. These observations, differ from those reported by Chen et al. with respect to the clinical impact of the infection[22]. While similarly observing the age-dependence of infection, the inoculations with SADS-CoV in their studies were lethal in the 2-day-olds, lethality decreasing in 5- and further in 7-day-old animals. A possible explanation for the difference might be the inoculation route, which was intragastrical in their studies and intraperitoneal in ours.

Chimeric mSADS-CoV exhibited subclinical infection in 7-day-old BALB/c mice, characterized by transient, low-level viral replication. However, in 2-day-old BALB/c mice the virus induced lethal infection, accompanied by significant growth retardation, elevated tissue viral titers, and mortality rates exceeding 60%. Postmortem examination revealed intestinal pathology characterized by wall edema, thinning, and the presence of foamy luminal contents, consistent with the SADS-CoV-induced phenomena observed in piglets [34] and in mouse models upon intracerebral infections [23, 34]. Although we have no data regarding the actual tropism of mSARS-CoV (nor of SADS-CoV) in the mice, the pathological changes caused by mSADS-CoV infection in 2-day-old BALB/c mice are very similar to those caused by SADS-CoV infection in porcine, the neonatal mouse model thus appearing to provide a convenient experimental model for studying SADS-CoV pathogenesis in swine. RDV, a broad-spectrum antiviral nucleotide analog, exerts its pharmacological effect through premature termination of viral RNA synthesis via inhibition of viral RdRp activity [35]. In vitro studies have established RDV’s inhibitory efficacy against multiple RNA viruses, encompassing SARS-CoV-2, SARS-CoV, MERS-CoV, Ebola virus (EBoV), respiratory syncytial virus (RSV), Nipah virus (NiV), porcine epidemic diarrhea virus (PEDV), porcine deltacoronavirus (PDCoV), and SADS-CoV [36–40]. The present study successfully implemented the mSADS-CoV mouse infection model for in vivo pharmacological evaluation. Unlike the non-treated mSADS-CoV-infected animals, most of which died, mice treated with RDV exhibited normal weight gain, no mortality, and a marked decrease in tissue viral load. Pathological lesions induced by viral infection were effectively alleviated, indicating that RDV potently inhibits mSADS-CoV replication in vivo, attenuates viral proliferation and infection across multiple tissues and organs, and consequently demonstrates significant antiviral activity. It is thus expected that the experimental platform will facilitate systematic screening of antiviral compounds targeting the viral replication machinery and will enable validation of host factors involved in viral replication processes.

Histopathological analysis of 2-day-old mSADS-CoV-infected mice at 5 dpi revealed extensive viral replication in multiple organs including pulmonary, intestinal, and splenic tissues, resulting in severe pathological alterations. The observed pulmonary and intestinal lesions exhibited remarkable similarity to those induced by SADS-CoV in neonatal mice [22]. Our immunohistochemical analysis demonstrated distinct tissue tropism patterns: anti-SADS-CoV N protein staining of mSADS-CoV-infected mouse tissues was predominantly localized in alveolar walls and intestinal basal layers. In general agreement herewith, upon intragastrical administration of SADS-CoV to 2-day-old mice viral antigen was found in alveolar walls, small intestinal villous epithelial cells and large intestinal basal layers [22], while intracerebral SADS-CoV administration to 7-day-old mice gave rise to viral antigen appearance in alveolar walls and intestinal epithelial cells [23]. These findings suggest that the spike gene modification of SADS-CoV that created the chimeric mSADS-CoV did not dramatically affect the overall tropism features of the virus in neonatal mice. SADS-CoV infection of pigs gives rise to severe and acute diarrhea and acute vomiting resulting in death but only in newborn piglets less than five days of age; older piglets and adult animals generally recover within a few days [34]. Histopathological evaluation of such lethally infected piglets revealed marked villus atrophy and viral antigen staining mainly in small intestine epithelial cells [34].

Despite its significant findings, this study has some methodological limitations. The mSADS-CoV model exhibits lethal infectivity exclusively in 2-day-old neonatal mice, which poses substantial experimental challenges due to their physiological immaturity, reduced robustness, and the heightened stress susceptibility of lactating dams. Furthermore, the evaluation of RDV’s *in vivo* efficacy was potentially compromised by suboptimal viral susceptibility, likely attributable to increased mouse stress responses induced by frequent post-administration manipulations, which may have influenced the viral infection process.

## Materials and methods

### Cell lines and viruses

African green monkey kidney Vero cells (ATCC CCL-81) were purchased from the American Type Culture Collection (ATCC), and mouse LR7 cells were kindly provided by Professor Chunhua Li from the Shanghai Academy of Agricultural Sciences. The SADS-CoV GDS04 strain (GenBank accession number: MF167434.1) was kindly provided by Professor Yongchang Cao from Sun Yat-sen University; MHV-A59 strain was purchased from the ATCC.

### Plasmid construction

Viral genomic RNA was isolated from the supernatants of SADS-CoV-infected Vero CCL-81 and MHV-A59-infected LR7 cells via TRIzol (Invitrogen). The extracted RNA was immediately used for cDNA synthesis according to the manufacturer’s instructions. Using the cDNA of these two viruses as templates, the required fragments were amplified, cloned and inserted into the pUC57 vector via 2× MultiF Seamless Assembly Mix (ABclonal) according to the manufacturer’s instructions (amplification primers are listed in Table 1). The resulting recombinant plasmids pUC57-SADS-ORF1ab-Spike-3′UTR and pUC57-SADS-ORF1ab-MHV Spike-3′UTR were confirmed by Sanger sequencing to ensure their accuracy and fidelity.

**Table 1.**
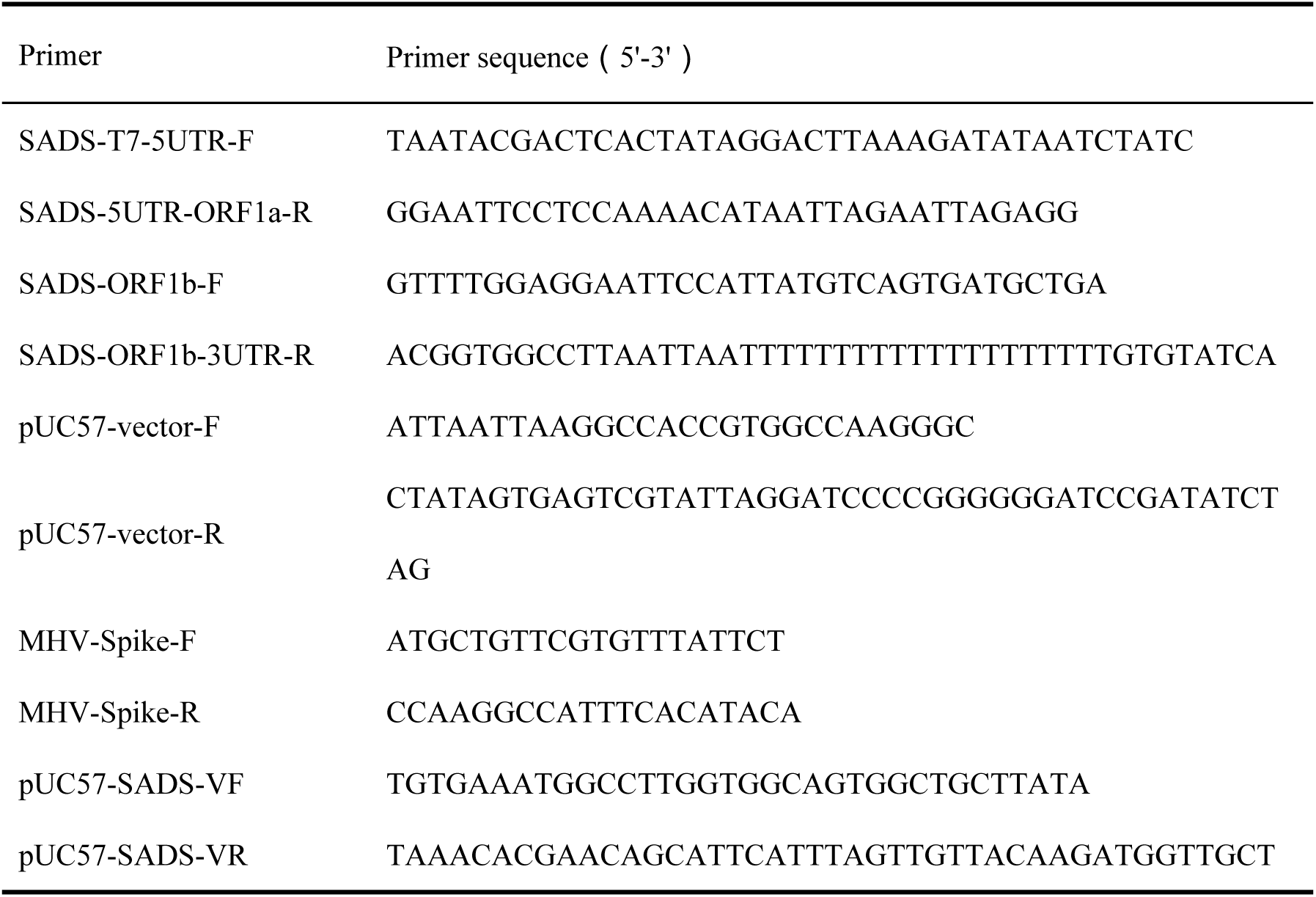
Primer Sequence List for Plasmid Construction.

### Rescue of the recombinant virus

The pUC57-SADS-ORF1ab-MHV Spike-3′UTR was linearized by PacI digestion and transcribed *in vitro* using an RNA transcription kit from Ambion following the manufacturer’s instructions. The resulting RNA transcripts were immediately placed on ice for electroporation. Vero CCL-81 cells that had been infected with SADS-CoV 5 hours earlier were trypsinized and washed twice with PBS. The SADS-CoV-infected Vero-CCL 81 cells were resuspended in PBS and the suspension was transferred to an electroporation cuvette. The *in vitro* transcribed RNA was gently admixed and the mixture was quickly placed in a Gene Pulser X cell electroporation system (Bio-Rad) for electroporation: a 450 V, 50 μF pulse was applied three times. The cells were quickly resuspended in cell growth medium and evenly plated onto a monolayer of LR7 cells. The cells were cultured overnight at 37 °C with 5% CO_2_ and washed with PBS, and the medium was replaced by maintenance medium (DMEM with 2% FBS) for continued culture, with regular observation for cytopathic effects.

### Immunofluorescence assay

The cell culture medium was discarded and the cells were washed three times with PBS before adding 200 μL of 4% formaldehyde and leaving the cells for 15 minutes at room temperature. The cells were washed three times with PBS, followed by the addition of 0.1% Triton X-100 solution for 15 minutes at room temperature for permeabilization. 3% BSA solution was added for blocking at room temperature for 1–2 hours. After three washes with PBS, the diluted primary antibody solution was added for incubation at room temperature for 2 hours. After washing three times with PBST, an Alexa Fluor 594/488-labeled donkey anti-mouse IgG secondary antibody dilution (1:500; Antgene, China) was added and the mixture was incubated at room temperature in the dark for 1 hour. DAPI (Solarbio, China) was added for incubation at room temperature in the dark for 10 minutes. The supernatant was discarded, and the cells were washed three times with PBS for fluorescence observation. The anti-SADS-CoV N protein monoclonal antibody, anti-SADS-CoV S protein monoclonal antibody, and monoclonal antibody targeting the MHV-S protein used in this study were prepared and preserved in our laboratory.

### Mouse infection experiments

3-week-old SPF-grade BALB/c and C57BL/6 mice were randomly divided into experimental and control groups, and inoculated with mSADS-CoV by intraperitoneal injection. Each mouse in the experimental group was injected with 1 mL of mSADS-CoV virus solution at a titer of 7×10^6.2^ PFU/mL, while the control group mice were injected with an equal volume of saline. 2-day-old and 7-day-old SPF-grade BALB/c mice were inoculated by intraperitoneal injection with 20 µL/g of an mSADS-CoV or SADS-CoV virus solution at a titer of 7×10^6.2^ PFU/mL; control animals received 20 µL/g of saline. This experimental protocol was approved by the Animal Ethics Committee of Huazhong Agricultural University, with ethics numbers HZAUMO-2024-0092 and HZAUMO-2024-0093.

### Evaluation of the *in vivo* inhibitory effect of RDV on viral infection in mice

2-day-old SPF-grade BALB/c mice were randomly divided into three groups, one virus infection group, one RDV-treated infection group and one control group. Due to the dependency of 2-day-old mice on maternal care during lactation, experimental groups were maintained in separate litters, with group sizes (5–8 pups) determined by natural birth rates. The mice in the two virus infection groups were inoculated by intraperitoneal injection with 20 µL/g of mSADS-CoV at a titer of 7×10^6.2^ PFU/mL. In parallel, mice in the control group similarly received saline. The mice in the RDV-treated infection group were administered RDV by subcutaneous injection of a dose of 25 mg/kg/d (drug solvent formulation: 2% DMSO, 40% PEG300, 5% Tween 80, 53% saline), the first administration at −1 dpi and subsequent additional administrations every 24 hours; these mice were designated the RDV-treated infection group. The animals in the mSADS-CoV group received subcutaneous injections with the same volume of drug solvent without the drug at the times when the RDV-treated infection group received the drug. At the same times, the animals in the mock group similarly received injections with the same volume of drug solvent. This experimental protocol was approved by the Animal Ethics Committee of Huazhong Agricultural University, with ethics number HZAUMO-2024-0094.

### Reverse transcription PCR and real-time quantitative PCR

After RNA was extracted from tissue samples, reverse transcription was performed according to the instructions of the Takara reverse transcription kit (RR036A). The primers used were designed on the basis of the conserved sequences of the SADS-CoV N gene and the MHV-A59 S gene (Table 2), and reverse transcription PCR was performed using DNA polymerase (CWBIO, China). The primers for quantitative fluorescence detection were designed on the basis of the conserved region of the SADS-CoV N gene, with the following primer sequences: forward primer SADS-CoV-qPCR-F: GTGCTAAAACTAGCCCCACAG; reverse primer SADS-CoV-qPCR-R: TGATTGCGAGAACGAGACTGTG. Fluorescence quantitative detection was performed via the Vazyme AceQ Universal SYBR qPCR Master Mix on the CFX96 and CFX384 instruments from Bio-Rad.

**Table 2.**
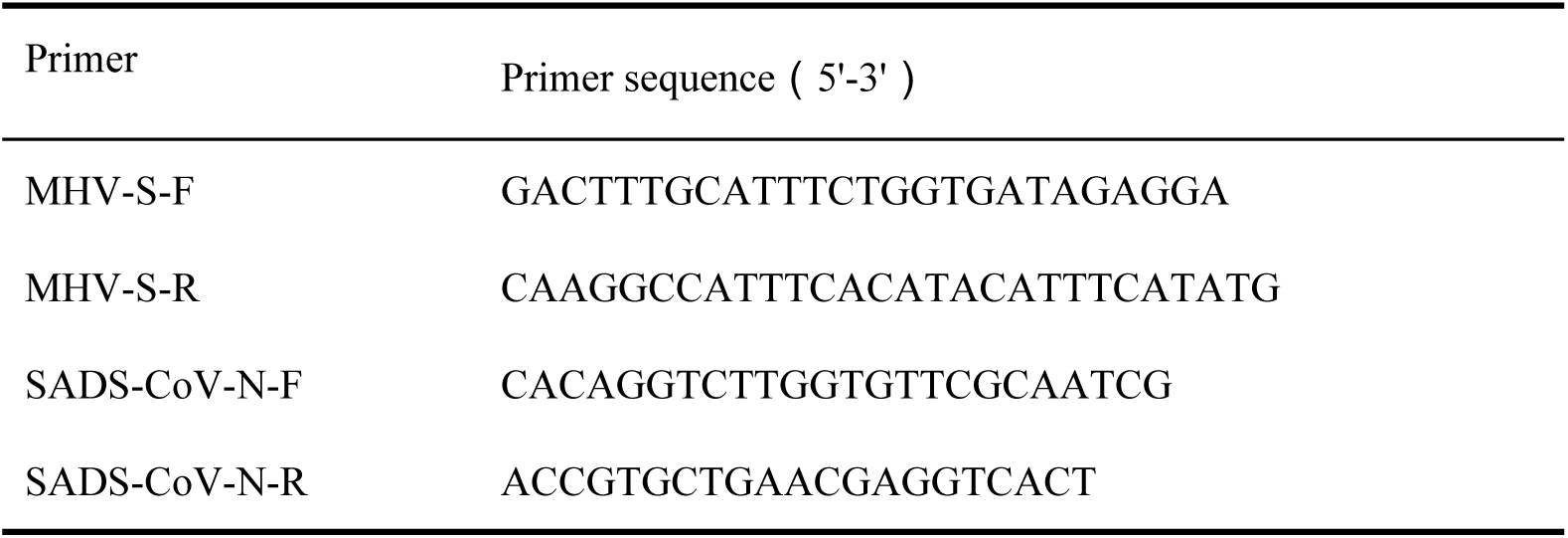
Primer sequence list for RT-PCR.

### Statistical analysis

Statistical analyses were performed via GraphPad PRISM 8.0.1. For comparisons between two experimental groups, two-tailed Student’s t tests were performed to determine significant differences. Statistical significance was as follows: *, *P*<0.05; **, *P*<0.01; ***, *P*<0.001; ****, *P*<0.0001; ns, not significant.

## Acknowledgments

The authors sincerely thank Professor Peter J.M. Rottier for his valuable guidance on manuscript preparation and insightful suggestions for revision. His expertise significantly enhanced the clarity and rigor of this work.

## Author Contributions

Conceptualization, Wentao Li, Mengjia Zhang, Hanyu Zhang; methodology, Wentao Li, Mengjia Zhang, Hongmei Zhu; investigation, Hanyu Zhang, Jiaru Zhou, Pengfei Li; formal analysis, Hanyu Zhang; validation, Mengdi Zhang, Ran Jing; resources, Wentao Li, Qigai He, Mengjia Zhang; supervision, Mengjia Zhang, Wentao Li; writing-original draft: Hanyu Zhang; writing-review and editing: Wentao Li, Mengjia Zhang, Yifei Lang, Qigai He and Hongmei Zhu; project administration, Wentao Li; funding acquisition, Wentao Li.

